# Graph embeddings for identifying symmetries in connectomes

**DOI:** 10.64898/2025.12.06.692615

**Authors:** Haozhe Shan, Ashok Litwin-Kumar

## Abstract

In many circuit models of neural computation, synaptic connections between neurons are organized according to their tuning to the variables being processed. The connectivities of canonical neural network models of head direction, spatial navigation, and orientation selectivity obey this principle and contain symmetries related to the angular and spatial variables they operate on. We develop a graph embedding algorithm to identify such symmetries. The algorithm segregates structure related to cell type from that related to the symmetries we seek to identify, distinguishing it from standard embedding methods. Our method successfully identifies rotational and translational symmetries in heading direction and visual projection neuron circuits using a connectome of the adult *Drosophila* brain, and it also identifies a toroidal symmetry in a synthetic connectome of grid cells in the medial entorhinal cortex. Such embedding geometries reveal the latent variables that are processed by a neural circuit and which cell types are responsible for this computation.

## 1 Introduction

Recently released synaptic connectome datasets detail the connections among up to 𝒪 (10^5^) neurons (Dorkenwald et al., 2024; Berg et al., 2025) and require scalable methods to infer the functions and computations performed by the neurons that comprise them. We propose a method motivated by the observation that symmetries in neural circuit connectivity are closely linked to the computations they perform (Langdon et al., 2023). For example, canonical ring attractor networks have recurrent weights with rotational symmetry (Fig. 1A left; Ben-Yishai et al., 1995), convolutional networks for visual processing have feedforward weights with translational symmetry (Fig. 1A center; Yamins and DiCarlo, 2016), and leading models of grid-cell systems have weights with a two-dimensional rotational symmetry (Fig. 1A right; Burak and Fiete, 2009). These symmetries arise from the shared coding of one or a set of low-dimensional variables. Locating neural populations whose connectivity obeys such symmetries may reveal circuits associated with computations involving these variables.

**Figure 1.**
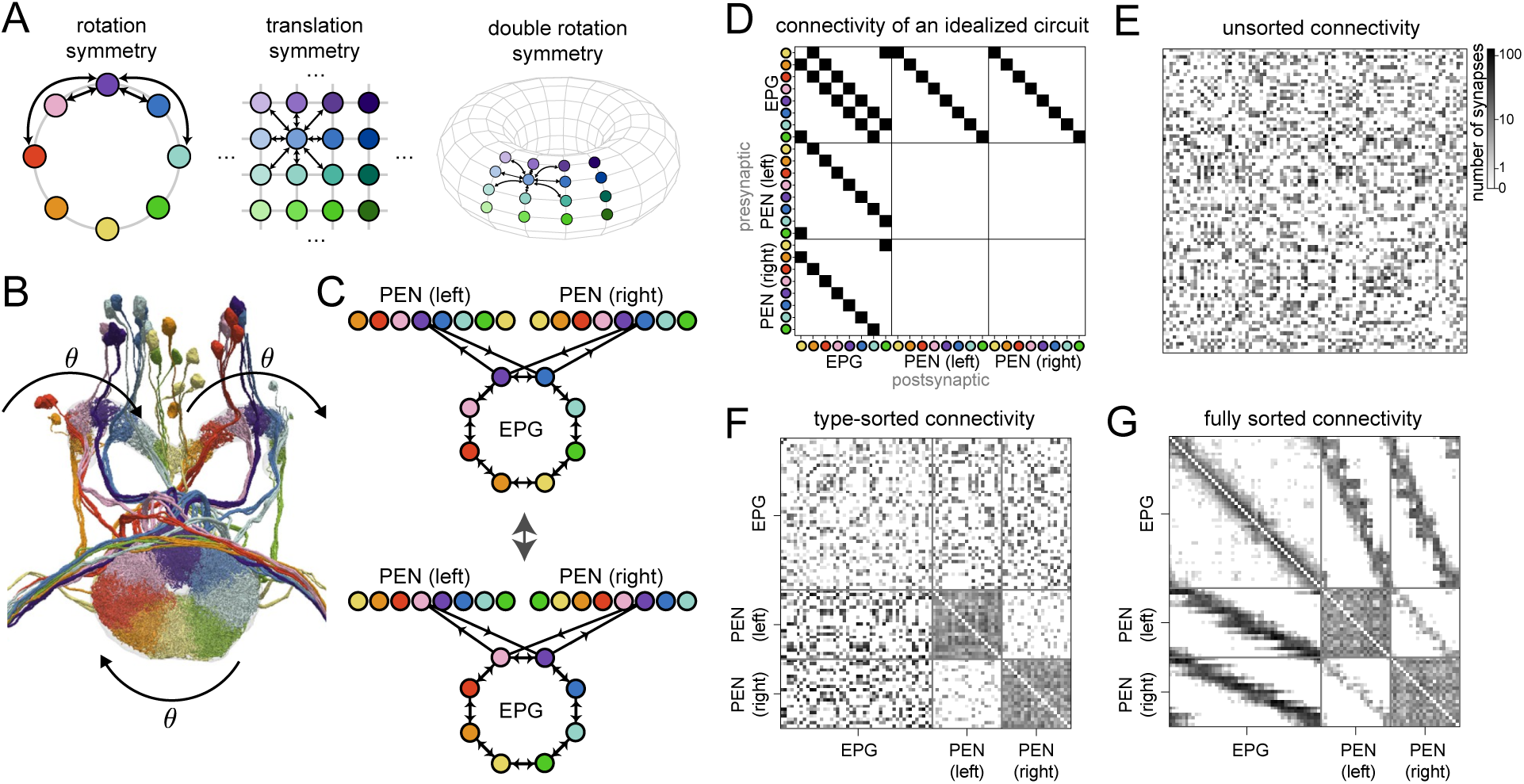
Symmetries and connectomes. **A** Schematics of different symmetries in the connectivities of neural circuit models. The hue and/or contrast indicate different values of the 1D or 2D variables associated with each neuron. **B** 3D reconstruction of a subset of neurons of the *Drosophila* central complex, adapted from Hulse et al. (2021). Colors identify neuron types and reflect organization with respect to an underlying angular variable *θ*. **C** Example of a graph automorphism. Top: Idealized model of EPG and PEN connectivity. For visual clarity, only one subset of connections between EPG and PEN neurons is shown. Node locations correspond to regions where neurons receive substantial dendritic input, rather than cell body locations. Bottom: Similar to top, but with neuron identity shifted with respect to *θ*. The resulting graph is isomorphic to the original (top). Connections between PEN neurons are omitted for simplicity. **D** Adjacency matrix of the idealized circuit illustrated in **C**. All the type-to-type submatrices are circulant. **E** Adjacency matrix of the EPG-PEN circuit in the hemibrain connectome. Without sorting, the matrix has a salt-and-pepper appearance that makes symmetries difficult to discern. All adjacency matrices are plotted on a log(1 + no. of synapses) scale. **F** Same as **E**, but with cells sorted by their cell types. **G** Same as **F**, but with cells sorted by their head-direction (*θ*) tuning. Contrast with idealized version in **D**. Type-to-type submatrices are not often not square, nor is any submatrix exactly circulant.

It is not clear how to efficiently search for this structure in connectome data, given that the symmetries that are present are likely to be approximate. The problem of detecting symmetries in graphs, for which the connectivity of certain nodes is perfectly equivalent to that of other nodes (formally, constructing a graph’s automorphism group), has been extensively studied. While no polynomial-time solution is known, practical algorithms exist (McKay and Piperno, 2014). However, connectome data contain potential variability due to both biological processes and imprecision in reconstruction. The problem of detecting “approximate” symmetries in noisy graphs has not been systematically examined.

To efficiently discover approximate symmetries (simply “symmetries” hereafter), we propose a method building on the graph embedding approach, where each node (neuron) is represented by a vector in a low-dimensional Euclidean space (Xu, 2021). Previous work has applied embedding methods to connectome analysis (e.g., Priebe et al., 2017; Pedigo et al., 2023; Matelsky et al., 2021; Mehta et al., 2023). However, due to the substantial heterogeneity in connectivity patterns across cell types, these methods tend to produce embeddings whose geometrical structure is dominated by cell type-level connectivity patterns (which cell type sends strong output to which other cell type, for example) while revealing little about the within-type connectivity that underlies computation. We overcome this problem by supplying the model with cell type information and explicitly parameterizing between-type connectivity statistics, thus segregating structure due to cell type and that due to underlying symmetry. We also develop algorithms for screening the learned embeddings for low-dimensional geometrical structures indicative of connectivity symmetries, such as rings.

Applied to a connectome of the adult *Drosophila* central brain (Scheffer et al., 2020), our method reliably identifies neurons with rotationally symmetric connections, such as EPG and Delta7 neurons. Additionally, it identifies a large number of cell types that make phase-modulated connections to the EPG circuit, suggesting that they are involved in heading direction-related computations. By searching for cell types with phase-shifted connections between them, we also identified circuits involved in angular velocity integration (Turner-Evans et al., 2017). We also applied our method to identify the retinotopic organization of visual projection neurons and how this information is transmitted to downstream neurons across the *Drosophila* brain. Finally, applying our method to a synthetic model of the connectivity of the medial entorhinal cortex, we demonstrate that it identifies neurons whose connectivity exhibits a 2D rotational symmetry, i.e., grid cells (Burak and Fiete, 2009).

## 2 Results

We begin by illustrating our approach using the connectome of the *Drosophila* central complex, whose neurons and connections are organized according to an underlying angular variable. We start by examining EPG and PEN neurons, which form contacts in the ellipsoid body and the left and right protocerebral bridge. Activity in these regions tracks heading direction (Seelig and Jayaraman, 2015; Green et al., 2017; Turner-Evans et al., 2017), and neurons can be labeled according to their preferred direction (Fig. 1B). EPG-EPG synapses are more numerous among neurons with similar preferred directions. EPG-PEN connectivity is organized similarly, but with an offset that enables PEN neurons to shift activity clockwise or anti-clockwise to integrate angular velocity (Green et al., 2017; Turner-Evans et al., 2017).

An idealized graph describing this circuit therefore contains edges connecting neurons with similar tuning (Fig. 1C, top). This graph has a nontrivial group of automorphisms corresponding to cyclic shifts of the neurons of each type (Fig. 1C, bottom), reflected in the block-circulant structure of its adjacency matrix (Fig. 1D).

We next compare this idealized graph to connectomic data (Scheffer et al., 2020). An unsorted connectivity matrix between EPG and PEN neurons obtained from the hemibrain connectome shows no discernible circulant structure (Fig. 1E). Sorting the rows and columns by cell types reveals cell-type-level structures but not the finer rotation symmetry (Fig. 1F). It is only after sorting individual neurons of each cell type by their preferred direction (*θ*) that block-diagonal bands indicative of circulant connectivity appear (Fig. 1G). This sorting was obtained by manual annotation guided by the tuning of central complex neurons being reflected in the physical locations of their processes (Hulse et al., 2021). Here, we ask how such structure could be extracted from connectivity and cell type information alone.

### 2.1 Graph embedding of synaptic connectivity data

Although the synaptic count data in Fig. 1G shows a clear circulant structure, it also displays variability. Unlike the idealized model (Fig. 1D), a graph with this adjacency matrix therefore contains no nontrivial automorphisms, ruling out approaches that search for automorphisms to identify such structure (McKay and Piperno, 2014). We therefore turned to graph embedding methods (Fig. 2A). Given a connectome of *N* neurons, we construct the adjacency matrix ***W*** ∈ ℤ^*N* ×*N*^, where *W*_*ij*_ is the estimated number of synapses from neuron *i* to neuron *j*. ***W*** therefore describes a weighted, directed graph. We aim to learn an embedding for each neuron *i*, which we denote by ***Z***_*i*_ ∈ ℝ^*D*^, from which ***W*** can be accurately reconstructed.

**Figure 2.**
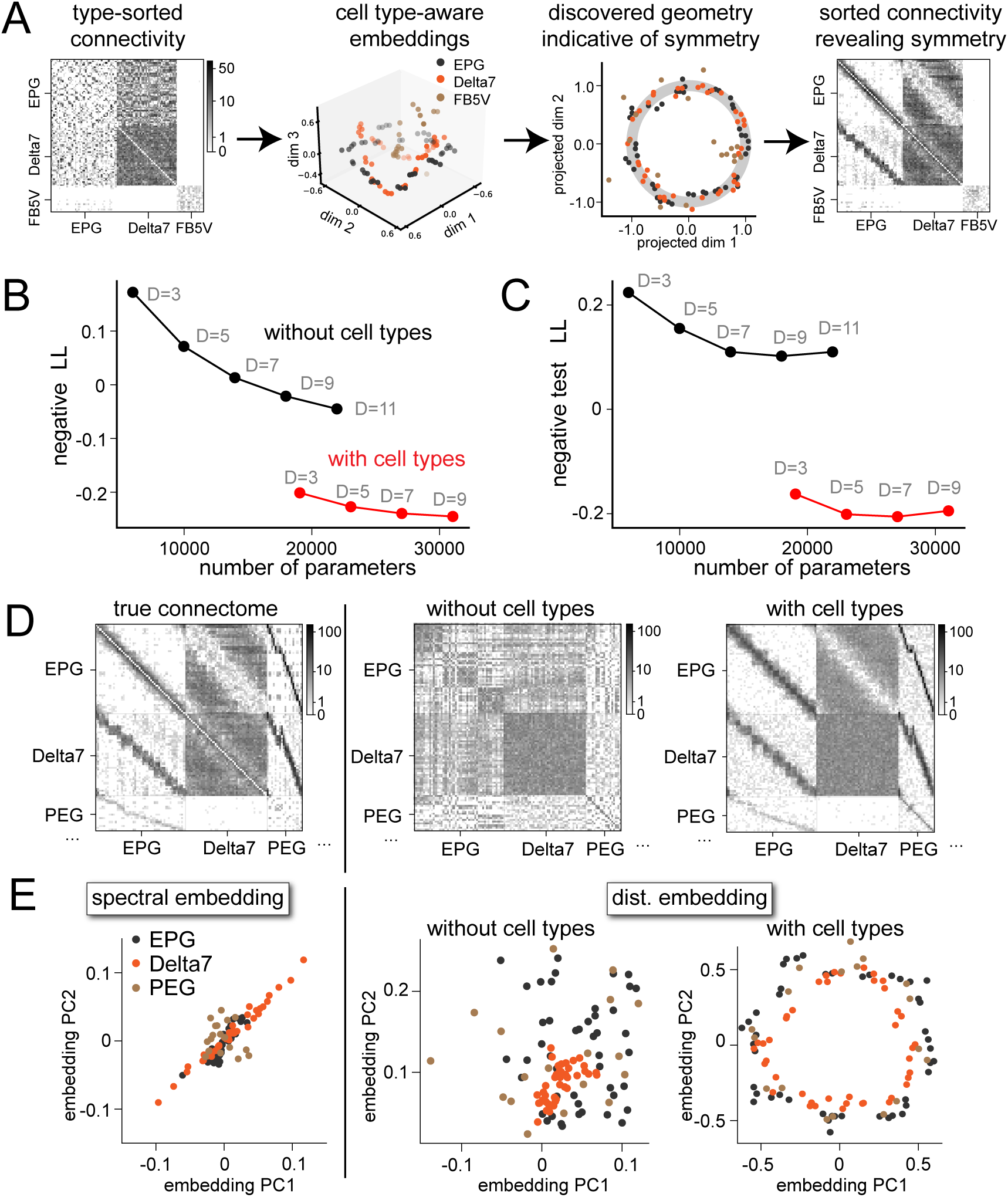
Embedding connectomes with cell type-specific parameters. **A** Overview of our method. The adjacency matrix from a connectome is first transformed into learned embeddings of neurons in a cell-type-aware manner. The embeddings are then screened for low-dimensional structures (circles in this example). Positions of neurons on such structures can be used to sort the adjacency matrix and reveal connectivity structures. **B** Negative log likelihood averaged over cell pairs of the central-complex adjacency matrix from models with and without cell-type-aware parameters. For reference, the raw data (adjacency matrix and cell-type indices) contain 4 × 10^6^ degrees of freedom. **C** Same as B, but for models trained on randomly chosen 80% of the cell pairs and tested on the 20% held-out pairs. **D** Left: the sorted ground-truth connectivity between Delta7, EPG and PEG neurons from hemibrain. Center: connectivity generated from a model without cell-type-specific parameters. Right: connectivity generated from a model with type-specific parameters. **E** Embeddings of EPG, Delta7, and PEG neurons using different methods. Left: spectral embeddings. Center: distance-based embeddings learned by a model without cell-type-specific parameters. Right: distance-based embeddings learned by a model with cell-type-specific parameters. Only the last set of embeddings show the three types of neurons as (approximately) concentric circles.

The central complex connectome displays structure both at the level of cell types and at the level of neurons’ angular tunings within a cell type (Fig. 1E-G). Connections among neurons of certain cell types are organized according to heading direction preference, but the specificity of this organization is type-specific. If the embeddings ***Z***_*i*_ and ***Z***_*j*_ solely reflected preferred heading direction, accurate reconstruction of *W*_*ij*_ would therefore require the mapping ***Z***_*i*_, ***Z***_*j*_ → *W*_*ij*_ to depend on the cell types to which neurons *i* and *j* belong.

Motivated by this observation, and by the fact that the cell types are often accurately inferred using morphological, connectivity, and other features (Schlegel et al., 2024), we study a graph embedding model that depends on neuron *i*’s cell type *c*_*i*_ ∈ {1, 2, … *N*_type_}, which we assume to be known (see Discussion). In this model, the weighted edge *W*_*ij*_ is drawn from an independent Poisson distribution with mean Ŵ_*ij*_, determined by an embedding function that depends on ***Z***_*i*_, ***Z***_*j*_, *c*_*i*_, and *c*_*j*_:

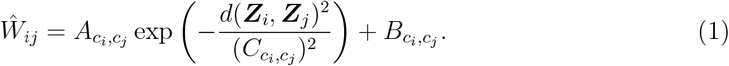

Here, *d*(***z, z***^*′*^) = ∥***z*** − ***z***∥^*′*^ is a distance function, taken to be Euclidean for now, and 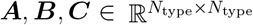 encodes cell-type-specific organizational structure. Specifically, *B*_*m*,*n*_ controls the average strengths of connections from cell type *m* to *n* regardless of the embeddings ***Z***, while *A*_*m*,*n*_ and *C*_*m*,*n*_ parameterize how the embeddings control connections between these cell types. To learn the embeddings, all trainable parameters are optimized to maximize the log likelihood log 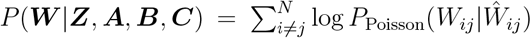, with a regularizer that penalizes large ***Z***. Altogether, the loss function to minimize reads

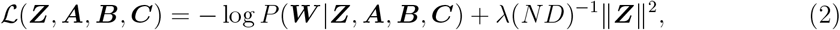

where *λ >* 0 is a hyperparameter (set to *λ* = 0.1). {***Z, A, B, C***} are optimized with Adam (Kingma and Ba, 2014), implemented in PyTorch. To prevent numerical issues while computing the Poisson likelihood, we passed all *Ŵ*_*ij*_ through the softplus function (softplus(*x*) = log(1 + *e*^*x*^)) to enforce nonnegativity.

We note that the model allows for limited asymmetry in the learned adjacency *Ŵ*. Connections between pairs of neurons belonging to the same type are assumed to be symmetric (up to asymmetries introduced by Poisson noise in the synaptic counts), but connectivity between neurons of different cell types can be asymmetric due to asymmetries in ***A, B, C***. We found these assumptions to be sufficient to identify much of the structure we describe here. Our models can be readily modified to accommodate full asymmetry by using the common approach of learning two *D*-dimensional embeddings for each neuron, a 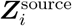 and a 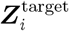, and computing *Ŵ* using 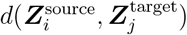 (Ou et al., 2016). We also describe a model that allows specific structured asymmetries within each cell type in Sec. 2.5.

### 2.2 Assessment of learned embeddings

We assessed the quality of the learned embeddings for the central complex connectome. For comparison, we also examined cell-type-unaware embeddings in which the parameters ***A, B, C*** are replaced with scalars that do not depend on cell type. For such models, ***Z*** must represent connectivity structure both at the level of types and within a type.

Including cell-type-specific parameters improves the model fit (Fig. 2B). A low-dimensional cell-type-aware embedding achieves a higher likelihood than a high-dimensional cell-type-unaware embedding, even when the total number of parameters is matched. This is not a result of overfitting, as cell-type-aware models trained on a randomly selected 80% of connections achieve a higher likelihood when assessed on their ability to predict a held-out 20% of connections (Fig. 2C). Cell-type-aware embeddings are therefore more accurate and require fewer dimensions to describe the central complex connectome.

The model described by Eq. 1 enables sampling from the learned distribution over adjacency matrices. A visual comparison between a ground-truth connectivity matrix and matrices generated by models with and without type-specific parameters is shown in Fig. 2D. Bands reflecting connectivity organized by angular tuning are better captured by the cell-type-aware embedding. Consistent with this, the top principal components of the cell-type-aware embedding vectors ***Z*** show a clear circular organization (Fig. 2E, right). This is not the case in the spectral embedding (left) nor in the embedding without cell type information (center). We conclude that segregating across-type and within-type level structures creates embeddings that more clearly portray the underlying organization with respect to heading direction.

### 2.3 Discovering connectivity structures by analyzing embedding geometry

The results of the previous sections demonstrate that symmetries in connectivity are reflected by geometrical structure in the learned embeddings. We developed a mathematical approach to identifying this structure, a problem that has two obstacles. First, we do not have access to the correct ordering of neurons with respect to the low-dimensional variables they encode (Fig. 3A), making our problem distinct from that of aligning neural representations for which this correspondence is assumed to be known (Kriegeskorte et al., 2008; Williams et al., 2021). Second, the geometry we seek may be present only in a particular projection of ***Z***, especially if there is structure in the connectivity beyond those variables we specify. The embeddings for the EPG neurons, for instance, contain structure in dimensions other than those that reflect their circular organization (Fig. 3B).

**Figure 3.**
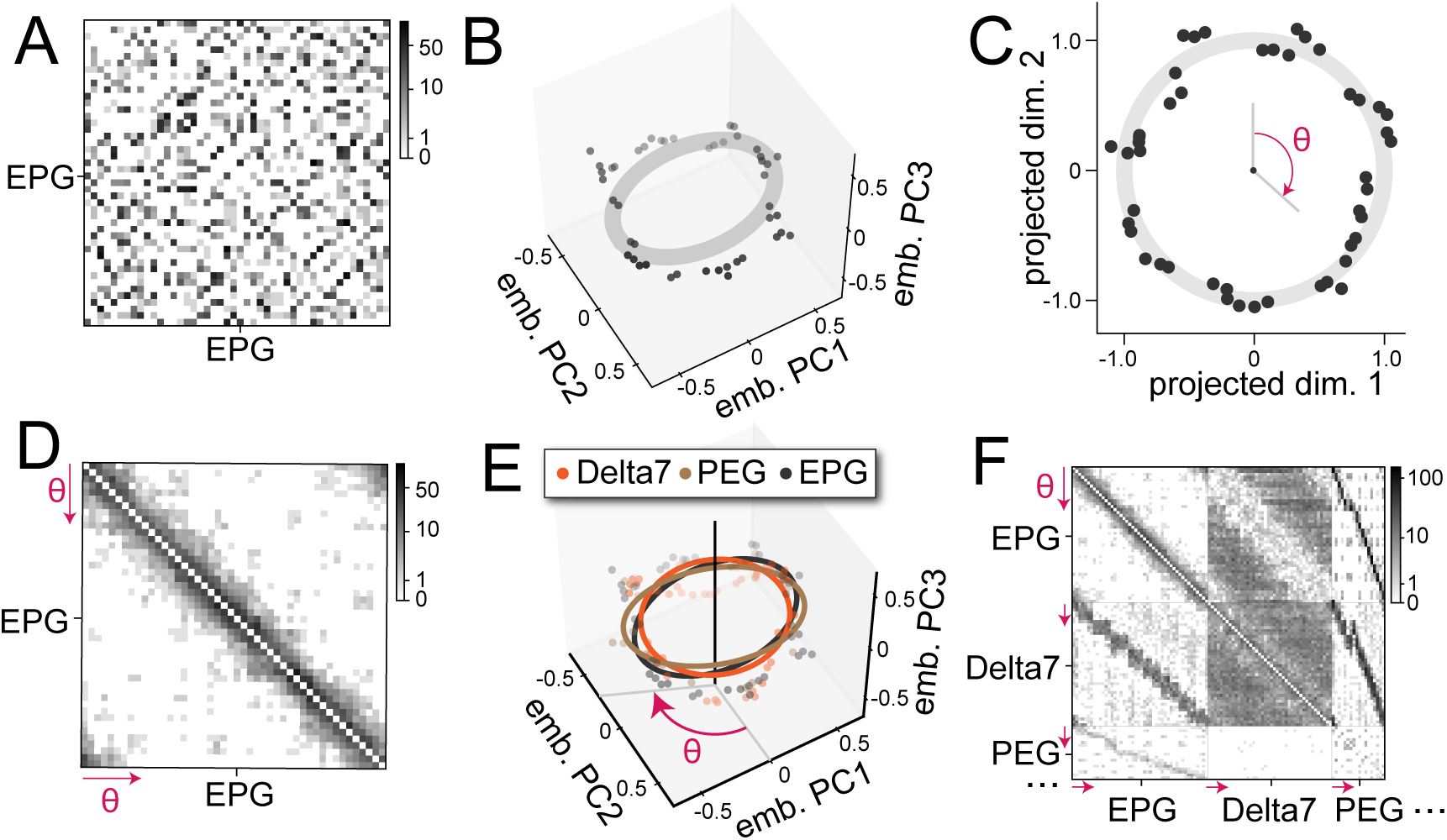
Identifying circles indicative of rotational symmetry in connectivity. **A** Unsorted connectivity between EPG neurons. **B** Top 3 principal components (PC) of the learned embeddings of EPG neurons. **C** EPG embeddings projected onto a circle using the circle-matching procedure. Individual neurons are assigned a *θ*. **D** Sorted connectivity between EPG neurons. The ordering is based on locations on the identified circle. **E** In the central complex, circles found in different cell types are approximately concentric. This structure allows the simultaneous sorting of connectivity between multiple cell types. **F** Simultaneous sorting reveals diagonal bands in between-type connectivity.

Overcoming these two obstacles amounts to finding a solution to the following optimization problem:

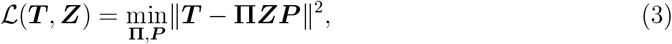

where ***T*** ∈ ℝ^*M* ×2^ is a “target” geometry, here a circle in 2D space 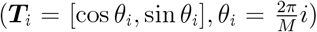. **Π** is optimized over *M* ×*M* permutation matrices and orders the neurons with respect to their tuning to *θ*. ***P*** is optimized over *D* × 2 real matrices and determines the embedding subspace in which a circular organization exists. ℒ (***T***, ***Z***) therefore measures the dissimilarity of the target geometry ***T*** and a projection of the reordered embeddings. Numerically, we solved Eq. 3 with a coordinate descent approach by alternating between optimizing **Π** using a fixed ***P***, a linear assignment problem (Burkard and Cela, 1999), and optimizing ***P*** under a fixed **Π**, a linear regression problem. The former subproblem can be solved with a convex relaxation (Jonker and Volgenant, 1988) and the latter has a closed-form solution, making the overall procedure efficient. This alternating optimization may suffer from local optima, but we found that this is easily overcome by repeating the procedure with 𝒪 (10) random initializations of **Π, *P***. We note that this procedure can be generalized to perform targeted identification of other low-dimensional structures by changing the target geometry ***T***, and in Sec. 2.7 we show how it can discover tori that indicate connectivity with two-dimensional rotational symmetry.

Applied to the embeddings of the EPG neurons, the projection and ordering procedure produces points distributed near the perimeter of a circle (Fig. 3C). Sorting neurons by their angle in this embedding subspace and reordering the rows and columns of the connectivity matrix accordingly recovers an approximately circulant matrix that reveals the rotational symmetry of the EPG neuron connectivity (Fig. 3D).

We note that embeddings for cell types whose recurrent connectivity is not tuned may nonetheless exhibit geometrical structure due to their interactions with other types. Sorting the connectivity correctly for these neurons critically depends on simultaneously embedding and sorting multiple cell types. Embedding vectors of neurons of different cell types that encode the same circular variable have approximately concentric embeddings (Fig. 3E). This allowed us to sort the connectivity between them in a way that is consistent across cell types (Fig. 3F). For example, although PEG neurons do not have structured recurrent connectivity among themselves, their structured projections to and from EPG and other neurons induce approximately circular embeddings (Fig. 3E) as well as diagonal bands in the corresponding blocks of the adjacency matrix (Fig. 3F). This example illustrates an important advantage of fitting shared embeddings across cell types rather than analyzing each type independently.

### 2.4 Screening for circular embeddings across central complex cell types

We next systematically searched for cell types involved in head-direction computation by screening for cell types with circular embeddings in the central complex. The adjacency matrix sorted by our method (Fig. 4A) contains numerous diagonal bands in blocks corresponding to across-type connections (Fig. 4B). To quantify the circularity of embeddings for each cell type, we defined a “circularity coefficient”, 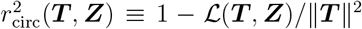. Analogous to the coefficient of determination, a value closer to 1 indicates that more variance after the projection can be explained by a circular organization.

**Figure 4.**
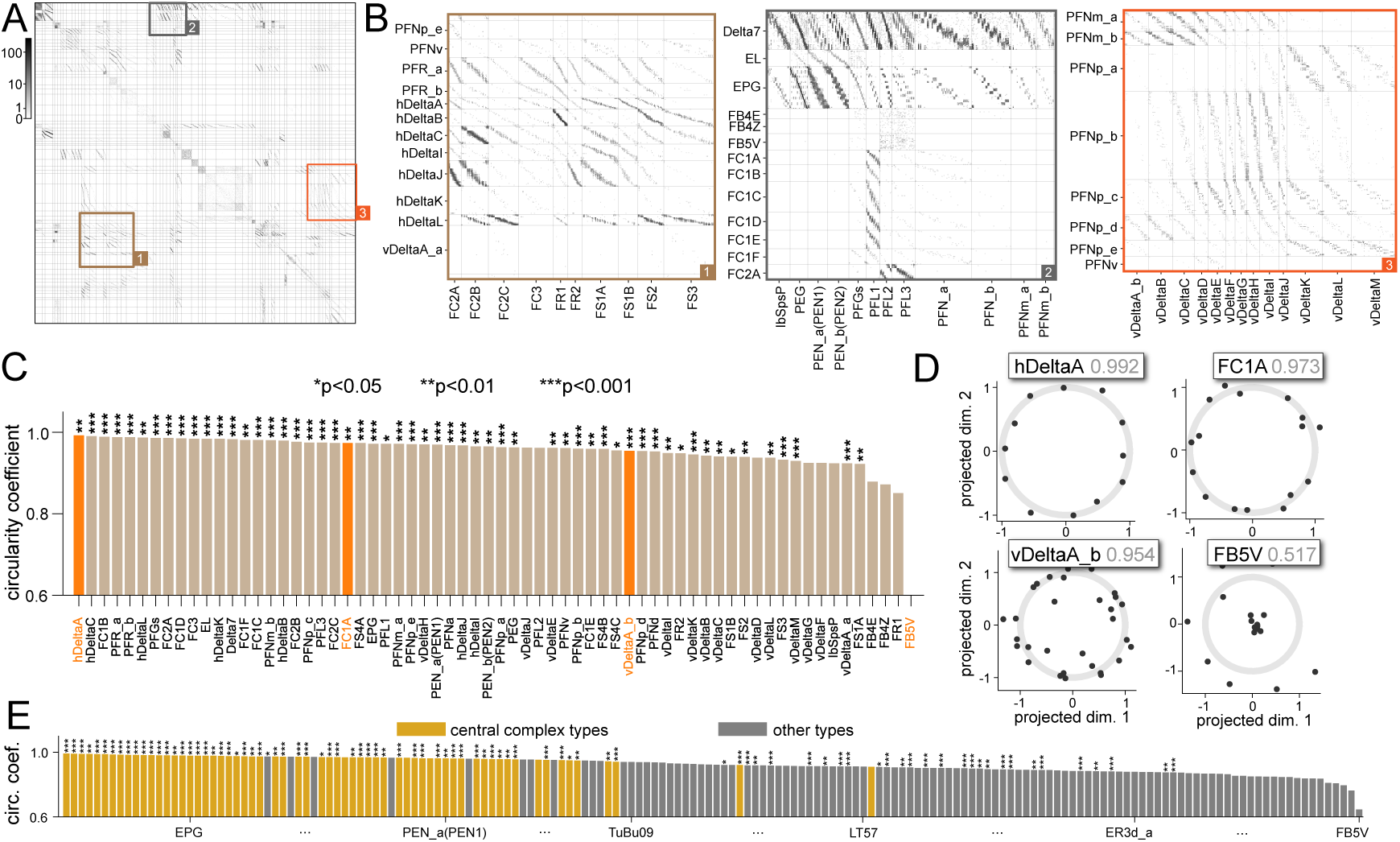
Identifying circles indicative of rotational symmetry in connectivity. **A** Adjacency matrix between neurons in the *Drosophila* central complex from the hemibrain dataset, after sorting using our method. **B** Sections of **A** enlarged for details. **C** Circularity coefficients (see text) for cell types in the central complex. Higher values indicate more circular embeddings for cells of this type. **D** Identified projections for example cell types (those shown with orange bars in **C**). In gray are the circularity coefficients. **E** Ranking cell types by their circularity coefficients in the full connectome. Cell types in the central complex are consistently found on top. Cell type labels are mostly omitted for brevity (See Fig. S1 for the fully labeled version; for central complex types, values may be different from those in **C** since it is generated using a different embedding).

To confirm that the identified structure is not a consequence of the freedom to choose an arbitrary ordering and projection, we computed a p-value for each cell type. We numerically estimated the probability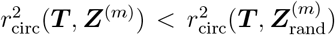, where the elements of 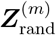 are i.i.d. and Gaussian, corresponding to an unstructured embedding (Methods 5.3). This analysis shows that the embeddings of the vast majority of central complex cell types have significantly circular embeddings (Fig. 4C,D; only types comprising more than 10 neurons are shown). Those with the lowest circularity coefficients are tangential neuron types (FB4E, FB4Z, FB5V). Tangential neurons innervate all columns of the fan-shaped body and thus discard heading direction information, consistent with this finding (Hulse et al., 2021).

Finally, to demonstrate that our approach does not require the subselection of a brain region containing a particular symmetry, we learned an embedding for every neuron in the hemibrain connectome simultaneously (Methods 5.1; Fig. S1A). As a quality check, we confirmed that the resultant embeddings of central complex neurons still exhibit the circular structures described in previous sections (Appendix Fig. S1B). In the connectome, cell types with high circular coefficients are overwhelmingly central complex types (Fig. 4E), consistent with their presumed role in navigation (Hulse et al., 2021).

### 2.5 Detecting phase-shifted connections

To perform certain heading direction computations, the *Drosophila* brain makes use of phase shifts: one cell type transmits a phase-shifted copy of its heading direction representation to another. A prominent example is the EPG-PEN circuit that performs angular-velocity integration (Turner-Evans et al., 2017). EPG-to-PEN weights and PEN-to-EPG weights are phase-shifted such that PEN activation “rotates” the activity bump that represents heading direction. This is accomplished by each PEN neuron receiving input from and projecting to EPG neurons with offset heading direction preferences (Fig. 5A). To detect such rotations based on connectome data, we modified our embedding method by including a type-specific rotation. We replaced the Euclidean distance in the embedding function (Eq. 1) with a new distance function:

**Figure 5.**
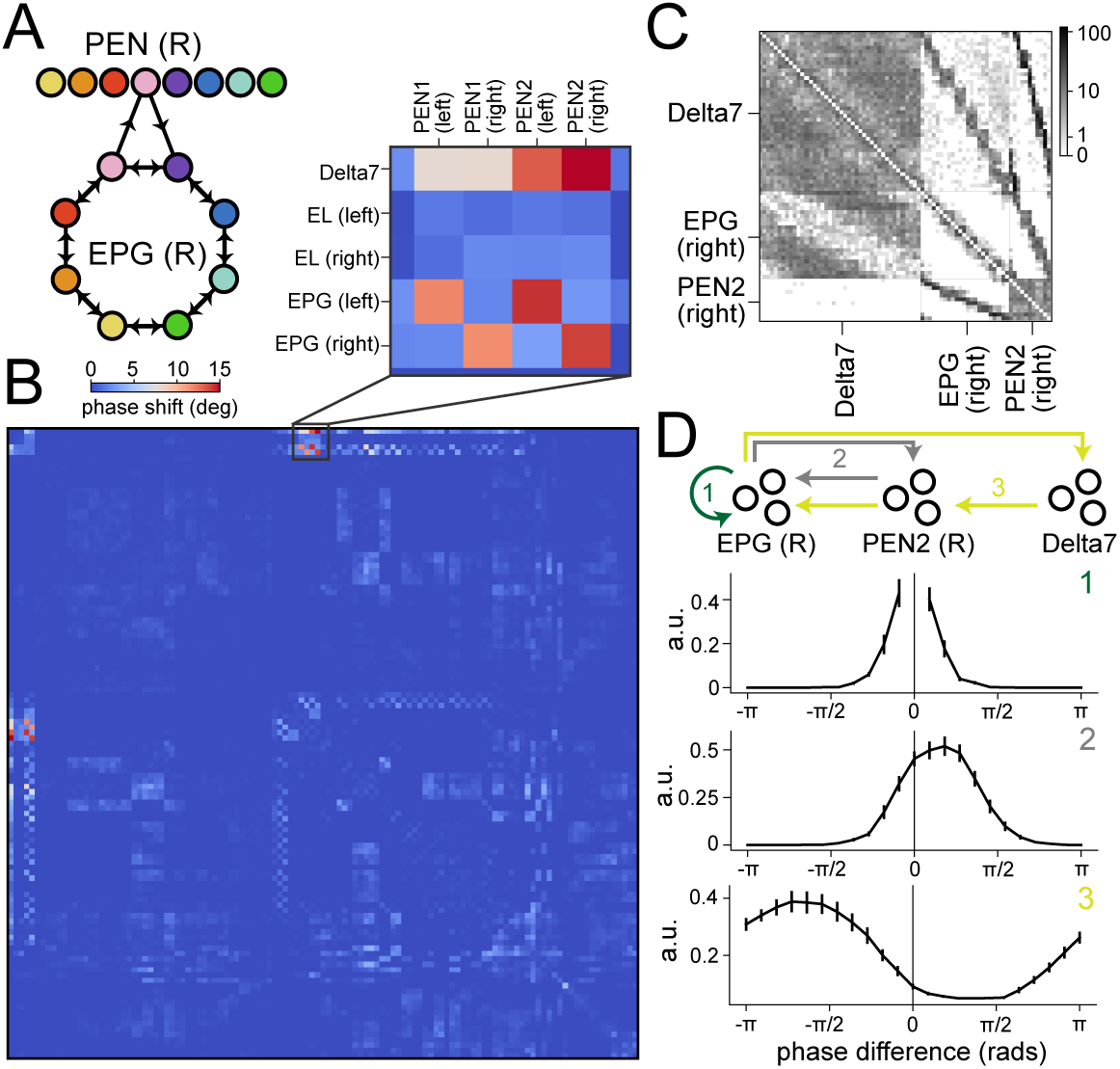
Detecting a phase shift connectivity structure. **A** Schematic of the phase shift structure between EPG and PEN neurons. **B** Detected phase shifts between pairs of cell types in the CX. The *m, n*th element shows 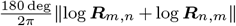, where {***R***_*m*,*n*_} are the unitary matrices in Eq. 4. A higher value indicates a larger relative rotation between the embeddings. Top inset: connections between Delta7, EPG (right) and PEN2 (right) neurons have significant relative rotations, indicating a phase shift structure. **C** Sorted connectivity between the three cell types. **D** Estimated connectivity tuning curves of EPG-to-EPG weights, EPG-to-PEN2-to-EPG effective weights and EPG-to-Delta7-to-PEN2-to-EPG effective weights. While the EPG-to-EPG weights have a symmetric (no phase shift) profile, the others are asymmetric.

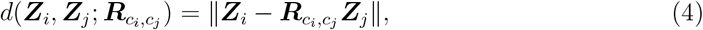

where 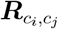is a *D* × *D* unitary matrix and is optimized analogously to ***A, B, C*** (see Methods and Appendix 6.3). Because phase shifts often differ between cells on the left and right sides of the brain, we also split the cells into left and right types. We analyze how much relative rotation is needed between cell types *m, n* by computing ∥log ***R***_*m*,*n*_ +log ***R***_*n*,*m*_∥, which reflects the magnitude of rotation by ***R***_*m*,*n*_***R***_*n*,*m*_.

This analysis provides an estimate of the relative rotation of embeddings across types (Fig. 5B). It suggests that the connectivity between EPG, Delta7, and PEN2 neurons (Fig. 5C) contains a phase shift structure. To confirm this, we estimated the connectivity tuning curves of EPG-to-EPG weights, EPG-to-PEN2-to-EPG effective weights, and EPG-to-Delta7-to-PEN2-to-EPG effective weights, where the effective weights are obtained by multiplying the corresponding adjacency matrices (Appendix 6.2). While recurrent connectivity among EPG neurons connects each neuron to others with similar phases, the connections through PEN2 and Delta7-PEN neurons peak at shifted phases (Fig. 5D), consistent with the presence of a phase shift structure. We conclude that, by appropriately parameterizing the embedding (Eq. 4), our method can identify both the geometry shared across cell types and the finer relationships between types, such as phase shifts.

### 2.6 Characterizing the retinotopic organization of *Drosophila* visual projection neuron connectivity

So far, we have illustrated our method by applying it to the central complex, a useful benchmark due to the substantial existing knowledge about its organization (Hulse et al., 2021). However, our general approach of searching for geometric structure in learned embeddings can be applied to other systems and symmetries. We next studied the structure of visual projection neurons (VPNs) in *Drosophila*, which receive feedforward visual input in the optic lobes and project to the central brain (Wu et al., 2016). Because these neurons’ axonal projections do not necessarily retain a retinotopic organization, it is not clear from morphology alone the extent to which this information is preserved in downstream projections.

We again analyzed an embedding of the entire hemibrain connectome (Methods 5.1) and focused the analysis on the embeddings of three groups of projection neurons: lobula columnar (LC) neurons, lobula-plate lobula columnar (LPLC) neurons, and medulla columnar (MC) neurons. Consisting of dozens of types, these neurons receive retinotopically organized input (Wu et al., 2016; Nern et al., 2025) and support distinct visually guided behaviors (Sen et al., 2017; Klapoetke et al., 2017; Ribeiro et al., 2018; Tanaka and Clark, 2022; Dombrovski et al., 2023). We first estimated putative receptive fields for each VPN based on anatomical information (Fig. 6A,B; Methods 5.2). We then asked whether the embeddings for each cell type, learned from connectivity data alone, reflect this retinotopy. We note that, because feedforward visual inputs to VPNs are not contained in the hemibrain connectome, these embeddings reflect only recurrent and output connectivity, and thus describe whether retinotopic information is preserved in such connections.

**Figure 6.**
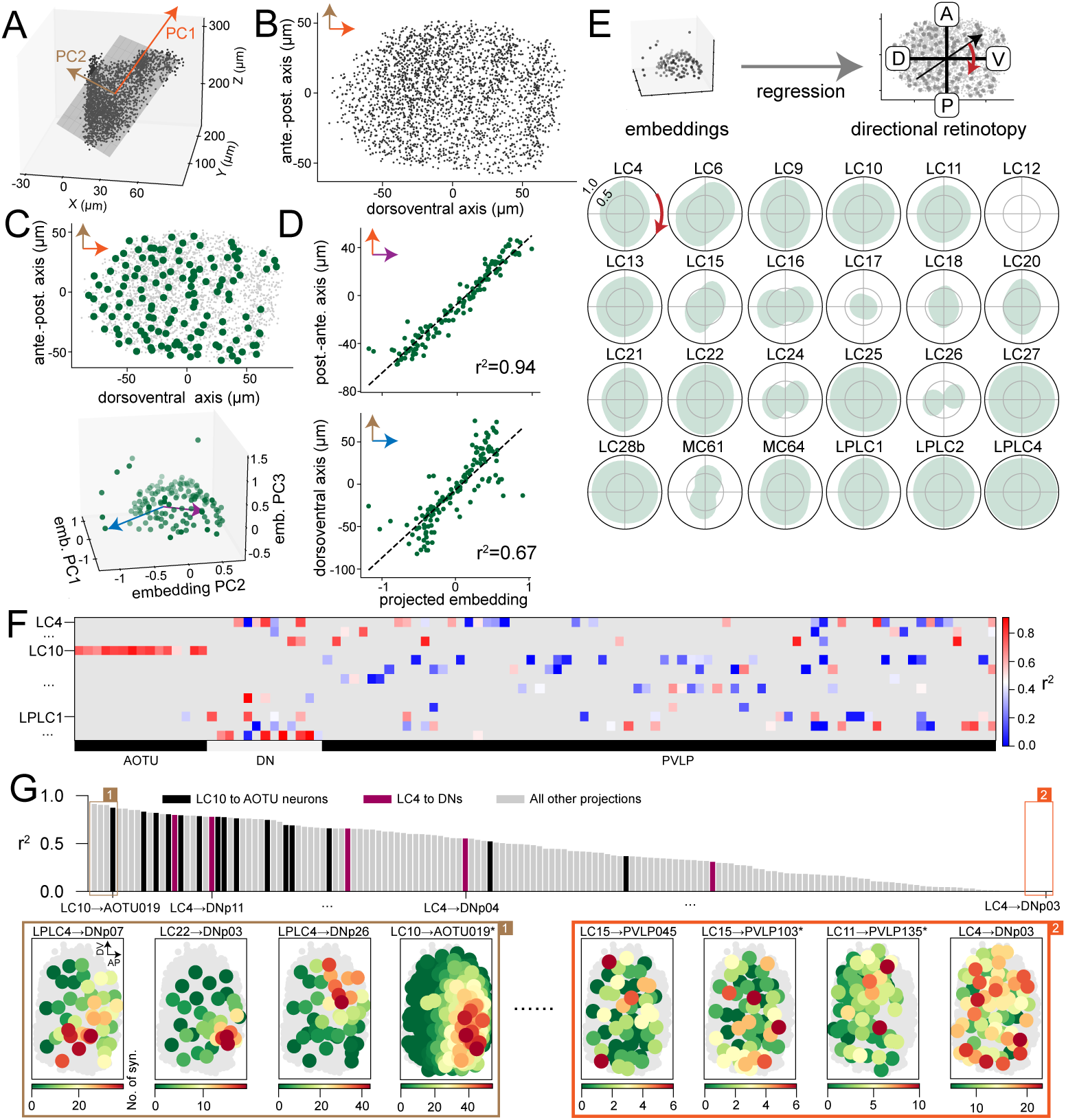
Detecting and quantifying retinotopy of VPNs and their downstream projections. **A** Centroids of the optic-lobe synapses of VPNs in 3D physical space. **B** The centroids after projection onto the top 2 PCs. The longer axis (PC1) corresponds to the dorsal-ventral axis of the visual field, while the shorter axis corresponds to the anterior-posterior axis. **C** Centroids (top) of LC9 neurons (green dots) are spread across the putative visual field. Embeddings of these neurons (bottom) also exhibit a 2D structure. **D** To quantify whether the embeddings reflect retinotopy, we performed linear regression from embeddings to the A-P or D-V coordinates of LC9 neurons. Brown and blue arrows indicate the best fitting directions in embedding space in **C**. **E** Directional retinotopy of all analyzed VPN types. A higher score along a direction indicates higher decodability of retinotopic coordinates along that direction from the embeddings. **F** Goodness of fit of projections from VPNs to their main downstream targets. The (*m, n*)-th element is the *r*^2^ between the model’s predicted numbers of synapses and the actual ones for projections from VPN type *m* and target type *n*. Gray areas correspond to projections with too few synapses (fewer than one synapse per pair of neurons on average). Only some example type labels are shown for legibility. **G** Ranking all the projections (non-gray elements in **F**). Some example projections are labeled. The retinotopy of the projection is visualized for the top 4 and bottom 4 projections. The visualizations are similar to **C** top, except now the number of synapses from each VPN to the target neuron is represented by the color. Most of these target types have one neuron per type; those with an asterisk have two each and an example one is shown. In the bar plot, some example projections are highlighted. Full information for **F** and **G** is available in Fig. S3.

Inspired by Dombrovski et al. (2023), we separately analyzed 1-D retinotopy along the dorsoventral (D-V) and anteroposterior (A-P) axes by regressing the D-V and A-P receptive field coordinates of each neuron against their embedding vectors (Fig. 6C). The degree to which the embeddings reflect retinotopic organization can then be quantified by the coefficients of determination (*r*^2^) of the fits (Fig. 6D). In the shown example, embeddings of the LC9 neurons are highly informative about the A-P coordinates and less so about the D-V coordinates, consistent with their axon topography (Dombrovski et al., 2023). We next generalized the 1D retinotopy analysis to non-cardinal directions (Fig. 6E). This analysis revealed that the connectivity of a broad range of LC/MC types exhibit reflects 2D retinotopy (e.g., LC10, LC28b), while for others this organization occurs primarily in one direction. Note that many of these cell types are known to have no axon topography, meaning that the organization we uncovered is not evident from their morphology (e.g., LC11, LC25; Dombrovski et al., 2023, 2025).

We next used our method to characterize how VPNs communicate to downstream targets, including neurons in the anterior optic tubercle (AOTU), the posterior ventrolateral protocerebrum (PVLP), and the motor descending neurons (DNs). Retinotopically specific connectivity, in which the number of input synapses a downstream neuron receives is enriched for VPNs that tile specific parts of the visual field, has been previously described in specific cases. These include selective projections from LC10 to AOTU neurons (Wu et al., 2016; Ribeiro et al., 2018) and “synaptic gradients” in connections such as those from LC4 to DNp02 and DNp11 neurons (Dombrovski et al., 2023).

Because many of these targets include cell types composed of few neurons, we learned a separate embedding of the connectome comprising all LC, LPLC, AOTU, PVLP neurons, and DNs, without a cutoff on the number of neurons per type. We also focused our ensuing analysis on VPNs with clear retinotopy in the embedding (Method 5.4). The degree to which a downstream target neuron’s VPN inputs are organized according to an underlying retinotopic variable can be quantified by the goodness-of-fit of our embedding model. We therefore computed the *r*^2^ of the model for each VPN-to-target projection with at least one synapse per pair of neurons on average and sorted the projections by this value (Fig. 6F,G; Methods 5.4). Projections known to be selective (e.g., LC10 to AOTU neurons; LC4 to DNp02 and DNp11 neurons) have high *r*^2^ values, whereas projections known to have no selectivity do not (e.g., LC4 to DNp03; Dombrovski et al., 2023). Visual inspection of these projections confirmed that the *r*^2^ value predicts retinotopic organization (Fig. 6G). Many projections that are highly selective have not been previously described. We conclude that the learned embedding model and its goodness-of-fit for different projections can be used as an efficient way to search for and summarize the retinotopic structure of VPN outputs. The analysis suggests that VPNs widely broadcast retinotopic information to their downstream targets. Detailed information about all analyzed projections is provided in Fig. S3.

### 2.7 Toroidal structure in a model of grid cells

Finally, we consider the toroidal symmetry in a circuit model of grid cells in the medial entorhinal cortex (MEC; Burak and Fiete, 2009). In this model, connections between grid cells are determined by their positions on a latent two-dimensional neural sheet with periodic boundary conditions. We investigated whether our approach can be used to discover this structure from the simulated connectome of such a circuit.

We first generated a grid cell circuit by modifying the connectivity structure in Burak and Fiete (2009) to obey Dale’s law (Methods 5.5). The circuit contains *n*_*E*_ × *n*_*E*_ excitatory neurons (i.e., the grid cells) but also *n*_*I*_ × *n*_*I*_ inhibitory neurons. Each neuron, excitatory or inhibitory, is defined by a location on the neural sheet, 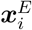 or 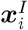, ranging from (− *n*_*E*_*/*2, −*n*_*E*_*/*2) to (*n*_*E*_*/*2, *n*_*E*_*/*2) (Fig. 7A, B). Neurons in the two populations are evenly distributed on this sheet. Weights between excitatory neurons are given by

**Figure 7.**
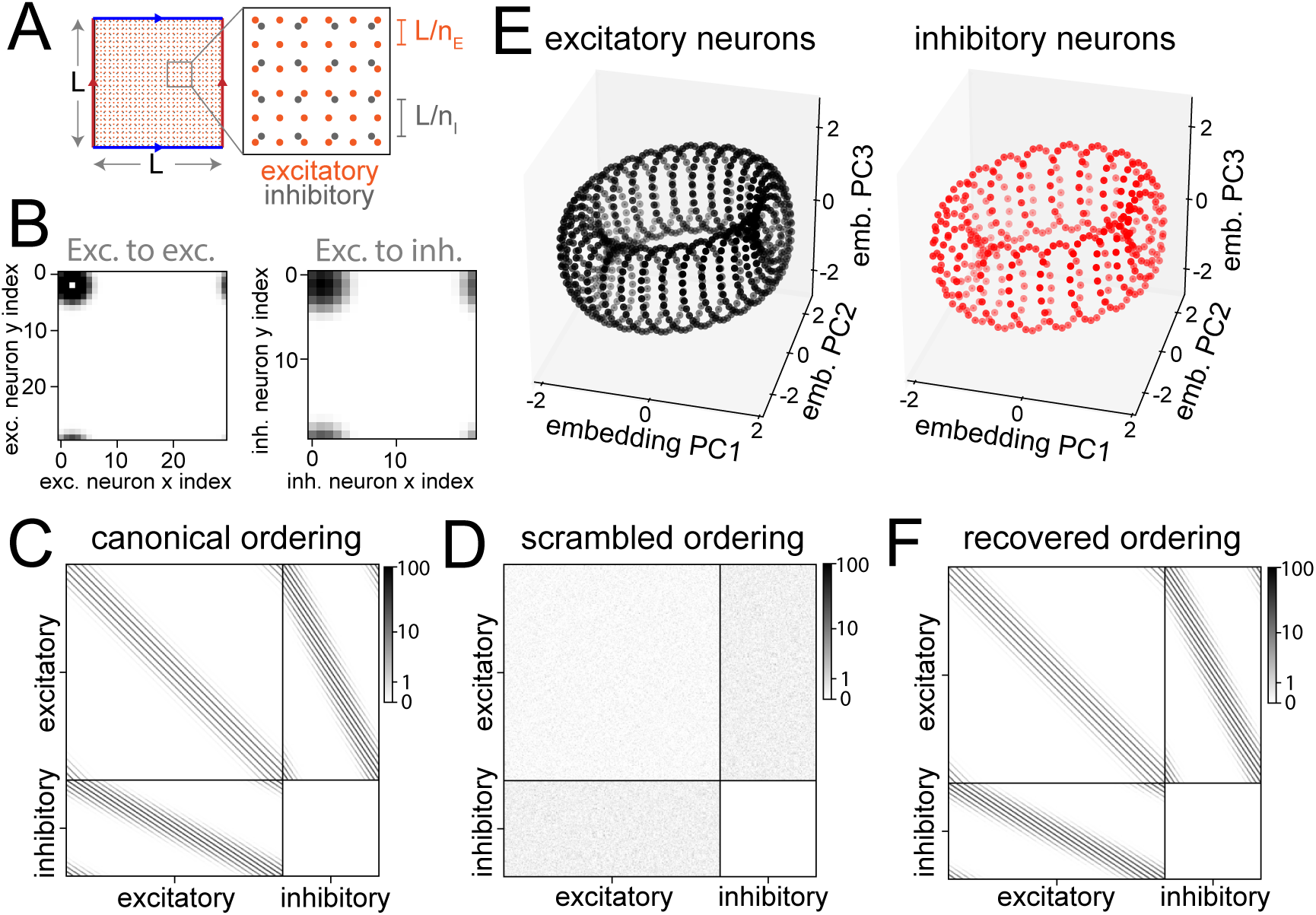
Detecting toroidal connectivity structure in a model of grid cells. **A** Excitatory and inhibitory neurons are placed on a 2D sheet with periodic boundaries. **B** Outgoing connectivity of an example excitatory neuron (coordinate (2, 2)) on the neural sheet in a grid cell model with 900 excitatory neurons and 400 inhibitory neurons. Synaptic strength of the example neuron onto excitatory neurons (top) and inhibitory neurons (bottom). **C** Ground-truth adjacency matrix of the model. Color scale represents connection strength. **D** Same as **C**, but with neuron ordering shuffled within each type. **E** Learned embeddings of excitatory neurons (left) form a Clifford torus (top 3 PCs are shown). Learned embeddings of inhibitory neurons (right) are distributed on the same torus but are spaced further apart. **F** Canonical ordering of neurons is recovered from the shuffled matrix in **D** using our approach.

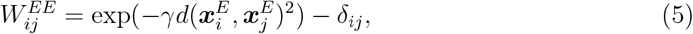

where *d*(***x, x***^*′*^) is the Euclidean distance under periodic boundaries and − δ_*ij*_ removes autapses. Weights from excitatory to inhibitory neurons and vice versa are given by

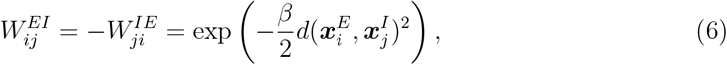

which generates an effective lateral inhibition among the excitatory neurons. We did not model recurrent inhibition. The parameters *γ* and *β* are estimated based on neural recordings (Burak and Fiete, 2009). We generated a simulated connectome with adjacency matrix *W*_*ij*_ ← *W*_0_|*W*_*ij*_|; *W*_0_ is chosen such that the maximum element is 100 (Fig. 7C, D). We interpret *W*_*ij*_ as the estimated strength of each connection, reflecting for instance presynaptic bouton or postsynaptic spine volume (Carlisle and Kennedy, 2005).

To accommodate the fact that *W*_*ij*_ is now continuous, we modified the likelihood function using a Gaussian distribution 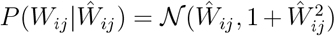; the choice of variance term allows greater variability for stronger synapses. We then learned embeddings for each neuron, treating excitatory and inhibitory neurons as two cell types and setting *D* = 5. The projections of both excitatory and inhibitory neurons appear as three-dimensional projections of a Clifford torus (Fig. 7E).

Similar to identifying an ordering for the ring attractor connectivity by matching neuronal embeddings to points on a circle, an ordering for this system can be obtained by matching to points on a Clifford torus. We solved the same optimization problem as in Eq. 3, except that ***T*** is replaced by an *N* × 4 Clifford torus and ***P*** is optimized over the space of *D* × 4 matrices. We first applied the method to the excitatory neurons and recovered a canonical ordering of them as well as a projection matrix ***P***_*E*_. We then applied the method to the inhibitory neurons but fixed the projection matrix to be ***P***_*E*_ to recover a canonical ordering of the inhibitory neurons. Reordering the excitatory and inhibitory neurons in this way recovered an ordered adjacency matrix (Fig. 7F).

## 3 Discussion

We have demonstrated that a cell-type-aware graph embedding method can efficiently identify connectivity structures that are organized according to low-dimensional variables. In the embedding space, identification of such organization amounts to describing the geometry of the neuronal embeddings. We presented an approach to this problem that involves matching points in the embedding space to a target geometry (Eq. 3), although other approaches to describe the topology or geometry of these points may also be effective (Carlsson, 2009). Our approach successfully identified the circular organization of the central complex connectome (Fig. 4), retinotopy in the organization of visual projection neurons (Fig. 6), and the toroidal organization of a simulated medial entorhinal cortex connectome (Fig. 7).

We have assumed that we have access to each neuron’s cell type prior to learning an embedding. This is motivated by the fact that existing methods can accurately identify cell types using morphological and connectivity features (Schlegel et al., 2024). It may be possible to use embeddings like those we describe here to facilitate cell type identification when such information is not available. Indeed, previous approaches have found embeddings that reflect such structure (Athreya et al., 2018). However, we have found that cell-type-unaware embeddings tend to be dominated by this structure, obscuring the symmetries that are the focus of our study (Fig. 2). Methods for simultaneous, demixed identification of cell types and the within-type symmetries we describe here are an interesting direction for future study. Seung (2009) described the problem of detecting underlying symmetries for synaptic chains and two-dimensional cognitive maps, and proposed a “graph layout” approach for the former problem. Other studies have used embeddings based on wiring length (Chen et al., 2006) or spectral methods (Petrovic et al., 2019) to analyze the *C. elegans* connectome. An analysis of the larval *Drosophila* mushroom body connectome using a dot-product embedding identified clusters related to distinct cell types (Athreya et al., 2018). An important distinction between these studies and our work is the cell type specificity of the mapping from embedding to connection probability, which we argue is beneficial for identifying the structure we seek.

We found that an embedding function based on Euclidean distance in the embedding space (Eq. 1) was sufficient to identify much of the structure present in the central complex connectome. We argue that this function should be cell-type-specific, but once this requirement is satisfied, the most effective parameterization of the function will likely depend on the system being studied. For example, we found that identifying asymmetries in the form of phase-shifted connections required an additional parameter (Eq. 4). Dot-product rather than distance-based embedding functions may also be appropriate, particularly for cell types organized based on multiple independent variables, which could correspond to distinct embedding subspaces. It may even be possible to learn embeddings in non-Euclidean spaces that better represent the structures we seek to identify (Bonnabel, 2013), or to incorporate graph convolution operations into the construction of the embedding (Kipf and Welling, 2017).

## 4 Acknowledgments

The authors would like to thank Larry Abbott for helpful feedback on the manuscript. This work was supported by the National Science Foundation and by DoD OUSD (R&E) under Cooperative Agreement PHY-2229929 (The NSF AI Institute for Artificial and Natural Intelligence), the Kavli Foundation, and the Gatsby Charitable Foundation (award #GAT3708). A.L.-K. was also supported by the National Science Foundation (award #2443158) and the National Institutes of Health (award #RF1DA060772).

## 5 Methods

### 5.1 Data and code

Most analyses presented here were performed on a subset of the hemibrain connectome of the *Drosophila* brain (Scheffer et al., 2020). The subset contains only cells in the central complex. In addition, we removed cell types with fewer than 10 neurons, since low-dimensional structures in the embedding space are difficult to discern with too few points. The list of cell types included in our data can be found in Appendix 6.1. This subset contains 1901 cells in 66 types. All ensuing analyses were done on embeddings fitted on this dataset, even if the embeddings/connectivity matrices of only a few cell types are shown in the figures. Results from fitting synthetic connectomes are discussed in Sec. 2.7. Unless otherwise noted, *D* = 5. ***A, B, C*** are initialized as uniform 1 matrices. ***Z*** is initialized as i.i.d. with 𝒩 (0, 1*/D*).

We also considered a full hemibrain embedding (Figs. 4, 6; Appendix Fig. S1), again with cell types having fewer than 10 neurons removed. This dataset contains 8145 neurons in 161 types. Hyperparameters and initialization methods are identical to those of the central-complex-only embedding.

Finally, in Sec. 2.6 we considered an embedding with all LC, LPLC, AOTU, and PVLP neurons and DNs. All of these cell types, regardless of the number of neurons per type, were included. This dataset contains 2894 neurons in 321 types. Again, hyperparameters and initialization methods are identical to those of the central-complex-only embedding.

### 5.2 Putative receptive fields for visual projection neurons

Visual projection neurons in the hemibrain connectome typically have recurrent synapses. Since the dendritic arbor of each visual projection neuron is known to be spatially localized (Wu et al., 2016), its relative location on the lobula can be taken as an estimate of its receptive field relative to the full visual field (Garner et al., 2024). For each neuron, we pooled all its synapses labeled as in the optic lobe and computed the centroid. Neurons without optic-lobe synapses (115 out of 2738) are excluded from subsequent analysis. Also, note that for MC neurons, input synapses from the lobula, but not from the medulla, are present in the connectome. MC62 neurons were excluded from further analysis because only 3 neurons had optic-lobe synapses. We then pooled these centroids for all neurons (Fig. 6A) and projected them into a two-dimensional plane obtained through principal components analysis, which collectively accounts for 94.6% of the variance. Based on the known organization of the lobula, we assumed the longer and shorter axes correspond to the dorsoventral (D-V) and anteroposterior (A-P) axes of the visual field, respectively (Fig. 6B; Garner et al., 2024; Nern et al., 2025). The location of each neuron’s centroid on the projection is taken as the center of its putative receptive field (Fig. 6B; Garner et al., 2024). Putative receptive fields of all columnar visual projection neurons analyzed are shown in Fig. S2. For all ensuing analyses, we excluded types for which we could not recover full-visual-field retinotopy (e.g., LC31). The types that were analyzed are listed in Fig. 6E.

### 5.3 Significance tests for circularity coefficients

To test the significance of a circularity coefficient *x*_0_ of *N* neurons in *D* dimensions, we generated 100 sets of *N* Gaussian i.i.d. vectors of *D* dimensions. For each set, we computed its circularity coefficient. We then generated a “null” distribution, 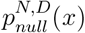 as a 1D Gaussian with mean and variance matching the 100 coefficients. The p-value of *x* is taken to be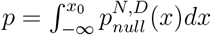.

### 5.4 Goodness of fit for projections

By “projection” from type *m* to type *n*, we refer to all the connections from the *N*_*m*_ type *m* neurons to the *N*_*n*_ type *n* neurons, summarized in a *N*_*m*_ × *N*_*n*_ submatrix, ***W*** ^*m*→*n*^, of the full adjacency matrix. Denote the model’s prediction for the block of interest as *Ŵ* ^*m*→*n*^. The goodness-of-fit of the model for this block is measured by the coefficient of determination,

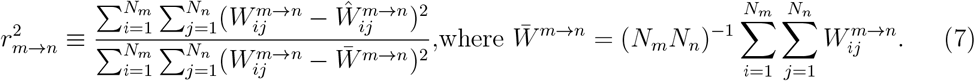

We first selected VPN types with good retinotopy in their embeddings (based on Fig. 6E; the criterion is the directional retinotopy in all directions must be greater than 0.5); : LC4, LC6, LC9, LC10, LC11, LC13, LC15, LC20, LC21, LC22, LC25, LC27, LC28b, LPLC1, LPLC2, and LPLC4. To guarantee sufficient statistical power, we excluded projections with fewer than one synapse per pair of neurons 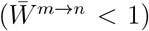. VPN types with all their projections excluded are not shown in Fig. 6F or Fig. S3.

### 5.5 Grid cell circuit with E/I neurons

The model consists of *n*_*E*_ × *n*_*E*_ excitatory neurons and *n*_*I*_ × *n*_*I*_ inhibitory neurons, posi-tioned on 2D square lattices with side length *n*_*E*_ and periodic boundary conditions (toroidal topology). Excitatory neurons are spaced at unit intervals, and inhibitory neurons have a spacing of *n*_*E*_*/n*_*I*_. Connectivity between two neurons is based on their geodesic distance on the surface, computed with the Euclidean distance metric.

The original model in Burak and Fiete (2009) contains only our excitatory neurons but modeled their connectivity as the sum of an excitatory component and an inhibitory component. Our E-to-E connectivity is the same as this excitatory component, while the E-to-I and I-to-E connectivity is such that multiplying the E-to-I and I-to-E submatrices results in an effectively inhibitory E-to-E connectivity that matches the inhibitory component in the original model.

For parameters *γ, β*, we followed the formula 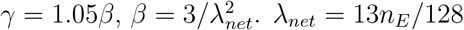.

### 6. Appendix

#### 6.1 Cell types considered to be in the central complex

We manually selected a subset of all cell types in the hemibrain dataset and considered these to be the “central complex” cells, based on the list in Scheffer et al. (2020), Appendix 1 Table 5. Note that cell types with fewer than 10 cells are always removed. The 66 types that remain are Delta7, EL, EPG, FB4E, FB4Z, FB5V, FC1A, FC1B, FC1C, FC1D, FC1E, FC1F, FC2A, FC2B, FC2C, FC3, FR1, FR2, FS1A, FS1B, FS2, FS3, FS4A, FS4B, FS4C, IbSpsP, PEG, PEN_a(PEN1), PEN_b(PEN2), PFGs, PFL1, PFL2, PFL3, PFNa, PFNd, PFNm_a, PFNm_b, PFNp_a, PFNp_b, PFNp_c, PFNp_d, PFNp_e, PFNv, PFR_a, PFR_b, hDeltaA, hDeltaB, hDeltaC, hDeltaI, hDeltaJ, hDeltaK, hDeltaL, vDeltaA a, vDeltaA b, vDeltaB, vDeltaC, vDeltaD, vDeltaE, vDeltaF, vDeltaG, vDeltaH, vDeltaI, vDeltaJ, vDeltaK, vDeltaL, and vDeltaM.

#### 6.2 Connectivity tuning curves

Let ***W***_sorted_ be the connectivity matrix, sorted by the estimated phases of neurons in each cell type (i.e., Fig. 2B). Let 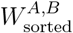 be a submatrix showing the weights from type A neurons and type B neurons. Then the A-to-B-to-A effective weights are defined as 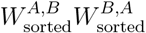 and similarly for A-to-B-to-C-to-A. As long as the chain begins and ends with type A, the result is a square matrix with rows and columns sorted by the estimated phases of type A neurons. The effective weights are normalized such that the maximum element is 1.

#### 6.3 Phase-shifted connections

If we label EPG neurons with evenly spaced angles 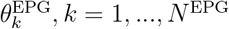 and PEN neurons with 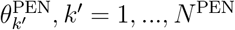, we can write the EPG-to-PEN weights as a function of 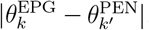. However, the PEN-to-EPG weights are “shifted” in that they are given by a function of 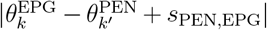, where *s*_PEN,EPG_ is the phase shift (schematized in Fig. 5A). We also note that this structure need not be between only two cell types and can be informally stated as the following: For cell types *m* = 1, 2, …, *M*, assign neurons from each type angles that are evenly distributed on the circle, 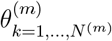 and suppose that the connectivity from any type *m* to another type *n* is predicted by a function of 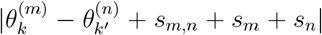, where {*s*_*m*,*n*_}, {*s*_*m*_} are scalar phases. A “phase shift structure” occurs when it is impossible to find a set of “absolute” phases {*s*_*m*_}_*m*=1,…,*M*_ such that ∀*m, n* : *s*_*m*,*n*_ = 0.

How might this structure be detected in a large connectome efficiently? We first make the following observation. For simplicity, suppose type A and type B neurons are equally numerous and respectively embedded on two circles. This implies that each neuron is assigned an angle (Fig. 5B). Suppose we assign angles to the neurons such that the type A neuron with angle 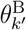 is connected to the type B neuron with angle 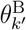, this implies that the two circles are aligned phase-wise (Fig. 5B, left). However, the type B → A weights suggest a different phase alignment, since the type A neuron with angle 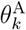 is now connected with the type B neuron with angle 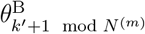 instead. In the embedding space, this means a relative rotation between the two circles, ***Z***^*A*^ and ***Z***^*B*^ (Fig. 5B). Thus, the phase shift structure between two cell types is reflected by a “tension” in the learned embeddings of these cells – the embeddings that best fit A → B weights and those that best fit B → A weights differ by a relative rotation between ***Z***^*A*^ and ***Z***^*B*^. Again, this can be stated in generality for multiple cell types. Suppose ***Z***^*m*=1,…,*M*^ are learned embeddings of *M* cell types. A phase shift structure occurs when connections from type *m* to type *n* are well predicted by 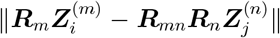, where {***R***_*m*_} and {***R***_*mn*_} are rotation matrices. Then a phase shift structure occurs when it is impossible to find {***R***_*m*_} such that ***R***_*mn*_ = 𝕀 for all *m, n*.

To numerically learn ***R***_*mn*_ using gradient descent on the loss function (Eq. 2), we parameterize it as 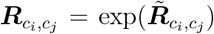, where exp is the matrix exponential and each 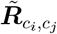 is a skew-symmetric matrix with *D*(*D* + 1)*/*2 free parameters to be optimized. Note that the rotations can be redundant in the sense that 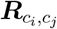 and 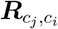may cancel each other. We thus add a term that penalizes the norm of 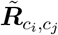to the loss function (Eq. 2; 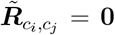results in 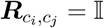).

**Figure S1:**
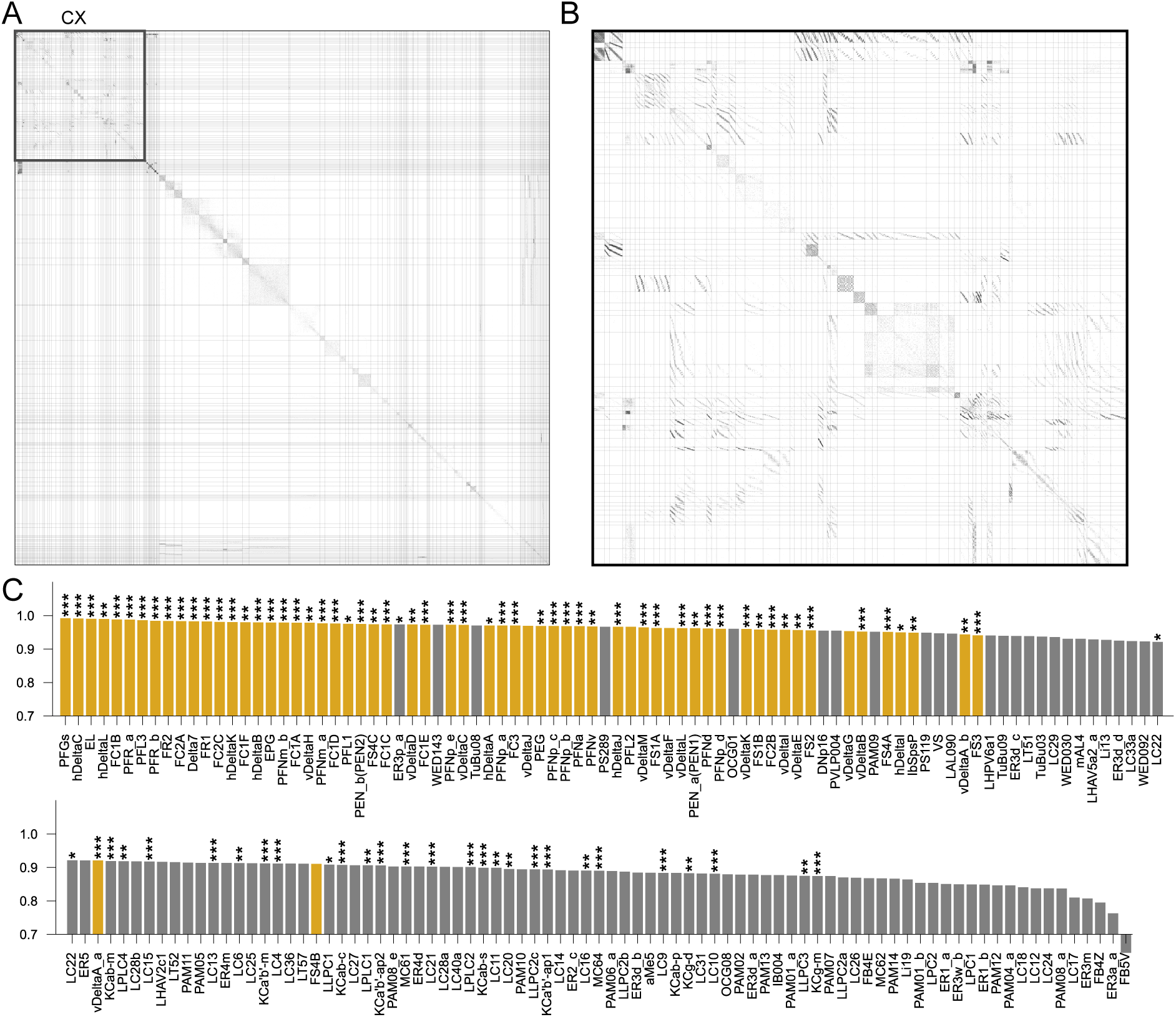
Embedding and sorting the entire hemibrain connectome simultaneously. **A** Sorted connectivity in the whole hemibrain connectome. **B** The section with connectivity within the central complex (CX), enlarged for details. Compare with Fig. 4A. **C** Circularity coefficient of cell types in the entire connectome. Same as Fig. 4E but with labels. \**p <* 0.05, \*\**p <* 0.01, \*\*\**p <* 0.001.

**Figure S2:**
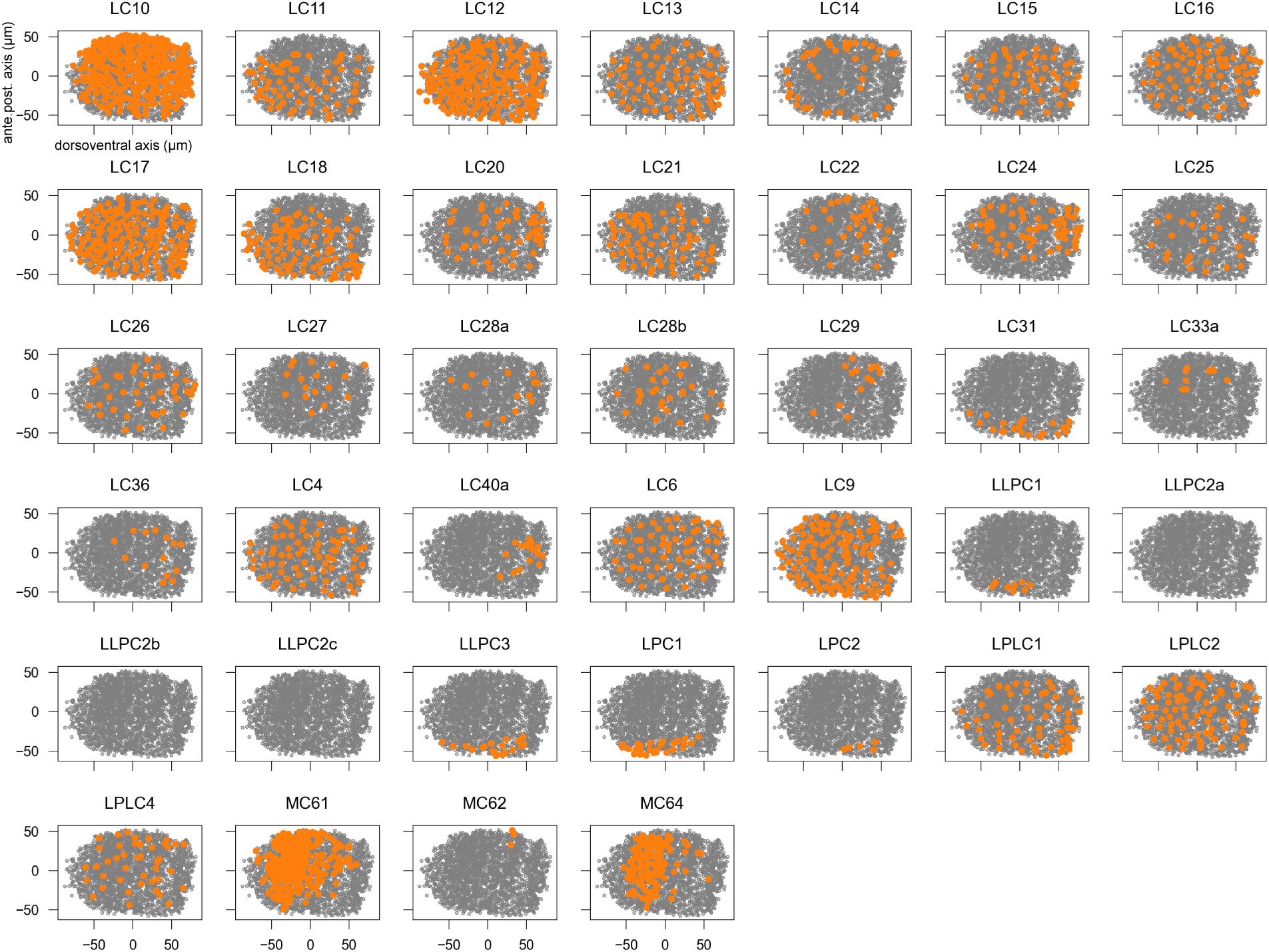
Estimated ground-truth retinotopy of columnar visual projection neurons. C.f. Fig. 6B. Each dot is the spatial centroid of optic-lobe synapses of a given neuron. Gray dots are all the cell types and orange dots are the given cell type. The centroids have been projected onto the top two PCs of all the centroids pooled together. See Fig. 6A.

**Figure S3:**
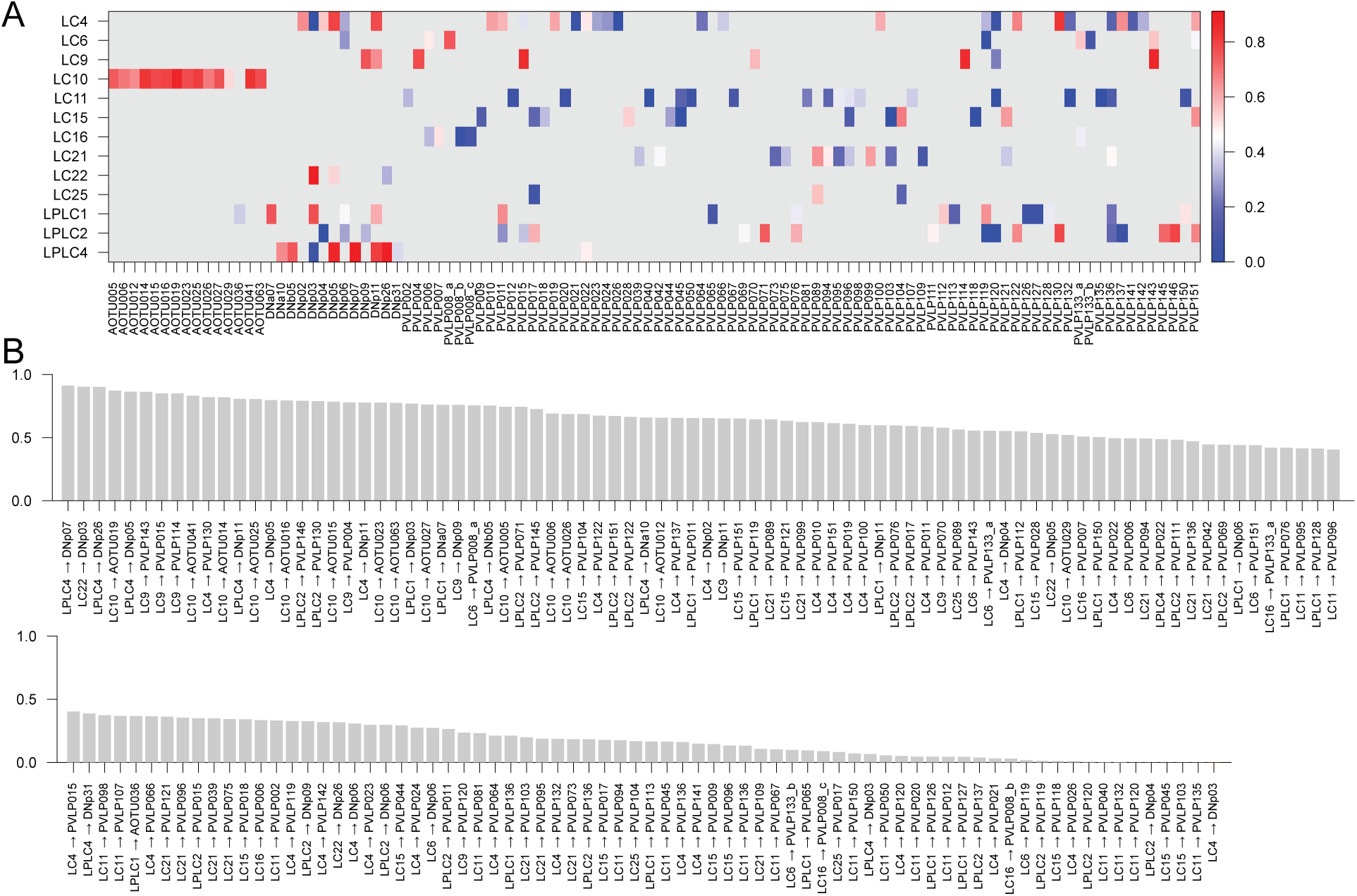
Model goodness-of-fit for projections from select VPNs to their target cell types. Same data as those presented in Fig. 6**F**,**G**. For each pair of cell types (types *m, n*), it is measured by computing the *r*^2^ between the actual numbers of synapses from each VPN neuron to each target neuron across all pairs (*N*_*m*_ × *N*_*n*_) and those predicted by the fitted model. Projections between cell types with less than, on average, one synapse per neuron pair were excluded from further analysis (Methods 5.4) The VPN types selected are those with high retinotopy scores, as displayed in Fig. 6E.

## References

A. Athreya, D. E. Fishkind, M. Tang, C. E. Priebe, Y. Park, J. T. Vogelstein, K. Levin, V. Lyzinski, Y. Qin, and D. L. Sussman. Statistical inference on random dot product graphs: a survey. Journal of Machine Learning Research, 18(226):1–92, 2018.

R. Ben-Yishai, R. L. Bar-Or, and H. Sompolinsky. Theory of orientation tuning in visual cortex. Proceedings of the National Academy of Sciences, 92(9):3844–3848, 1995.

S. Berg, I. R. Beckett, M. Costa, P. Schlegel, M. Januszewski, E. C. Marin, A. Nern, S. Preibisch, W. Qiu, S.-y. Takemura, et al. Sexual dimorphism in the complete connectome of the drosophila male central nervous system. bioRxiv, pages 2025–10, 2025.

S. Bonnabel. Stochastic gradient descent on riemannian manifolds. IEEE Transactions on Automatic Control, 58(9):2217–2229, 2013.

Y. Burak and I. R. Fiete. Accurate path integration in continuous attractor network models of grid cells. PLoS computational biology, 5(2):e1000291, 2009.

R. E. Burkard and E. Cela. Linear assignment problems and extensions. In Handbook of combinatorial optimization: Supplement volume A, pages 75–149. Springer, 1999.

H. J. Carlisle and M. B. Kennedy. Spine architecture and synaptic plasticity. Trends in neurosciences, 28(4):182–187, 2005.

G. Carlsson. Topology and data. Bulletin of the American Mathematical Society, 46(2):255–308, 2009.

B. L. Chen, D. H. Hall, and D. B. Chklovskii. Wiring optimization can relate neuronal structure and function. Proceedings of the National Academy of Sciences, 103(12):4723– 4728, 2006.

M. Dombrovski, M. Y. Peek, J.-Y. Park, A. Vaccari, M. Sumathipala, C. Morrow, P. Breads, A. Zhao, Y. Z. Kurmangaliyev, P. Sanfilippo, et al. Synaptic gradients transform object location to action. Nature, 613(7944):534–542, 2023.

M. Dombrovski, Y. Zang, G. Frighetto, A. Vaccari, H. Jang, P. S. Mirshahidi, F. Xie, P. Sanfilippo, B. W. Hina, A. Rehan, et al. Molecular gradients shape synaptic specificity of a visuomotor transformation. Nature, pages 1–10, 2025.

S. Dorkenwald, A. Matsliah, A. R. Sterling, P. Schlegel, S.-C. Yu, C. E. McKellar, A. Lin, M. Costa, K. Eichler, Y. Yin, et al. Neuronal wiring diagram of an adult brain. Nature, 634(8032):124–138, 2024.

D. Garner, E. Kind, J. Y. H. Lai, A. Nern, A. Zhao, L. Houghton, G. Sancer, T. Wolff, G. M. Rubin, M. F. Wernet, et al. Connectomic reconstruction predicts visual features used for navigation. Nature, 634(8032):181–190, 2024.

J. Green, A. Adachi, K. K. Shah, J. D. Hirokawa, P. S. Magani, and G. Maimon. A neural circuit architecture for angular integration in drosophila. Nature, 546(7656):101–106, 2017.

B. K. Hulse, H. Haberkern, R. Franconville, D. B. Turner-Evans, S.-y. Takemura, T. Wolff, M. Noorman, M. Dreher, C. Dan, R. Parekh, et al. A connectome of the drosophila central complex reveals network motifs suitable for flexible navigation and context-dependent action selection. Elife, 10, 2021.

R. Jonker and T. Volgenant. A shortest augmenting path algorithm for dense and sparse linear assignment problems. In DGOR/NSOR: Papers of the 16th Annual Meeting of DGOR in Cooperation with NSOR/Vorträge der 16. Jahrestagung der DGOR zusammen mit der NSOR, pages 622–622. Springer, 1988.

D. P. Kingma and J. Ba. Adam: A method for stochastic optimization. arXiv preprint 1412.6980, 2014.

T. N. Kipf and M. Welling. Semi-supervised classification with graph convolutional networks. In International Conference on Learning Representations, 2017. URL https://openreview.net/forum?id=SJU4ayYgl.

N. C. Klapoetke, A. Nern, M. Y. Peek, E. M. Rogers, P. Breads, G. M. Rubin, M. B. Reiser, and G. M. Card. Ultra-selective looming detection from radial motion opponency. Nature, 551(7679):237–241, 2017.

N. Kriegeskorte, M. Mur, and P. A. Bandettini. Representational similarity analysis-connecting the branches of systems neuroscience. Frontiers in systems neuroscience, 2:249, 2008.

C. Langdon, M. Genkin, and T. A. Engel. A unifying perspective on neural manifolds and circuits for cognition. Nature Reviews Neuroscience, 24(6):363–377, 2023.

J. K. Matelsky, E. P. Reilly, E. C. Johnson, J. Stiso, D. S. Bassett, B. A. Wester, and W. Gray-Roncal. Dotmotif: an open-source tool for connectome subgraph isomorphism search and graph queries. Scientific Reports, 11(1):13045, 2021.

B. D. McKay and A. Piperno. Practical graph isomorphism, ii. Journal of symbolic computation, 60:94–112, 2014.

K. Mehta, R. F. Goldin, and G. A. Ascoli. Circuit analysis of the drosophila brain using connectivity-based neuronal classification reveals organization of key communication pathways. Network Neuroscience, 7(1):269–298, 2023.

A. Nern, F. Loesche, S.-y. Takemura, L. E. Burnett, M. Dreher, E. Gruntman, J. Hoeller, G. B. Huang, M. Januszewski, N. C. Klapoetke, et al. Connectome-driven neural inventory of a complete visual system. Nature, pages 1–13, 2025.

M. Ou, P. Cui, J. Pei, Z. Zhang, and W. Zhu. Asymmetric transitivity preserving graph embedding. In Proceedings of the 22nd ACM SIGKDD international conference on Knowledge discovery and data mining, pages 1105–1114, 2016.

B. D. Pedigo, M. Powell, E. W. Bridgeford, M. Winding, C. E. Priebe, and J. T. Vogelstein. Generative network modeling reveals quantitative definitions of bilateral symmetry exhibited by a whole insect brain connectome. Elife, 12:e83739, 2023.

M. Petrovic, T. A. Bolton, M. G. Preti, R. Liégeois, and D. Van De Ville. Guided graph spectral embedding: Application to the c. elegans connectome. Network Neuroscience, 3(3):807–826, 2019.

C. E. Priebe, Y. Park, M. Tang, A. Athreya, V. Lyzinski, J. T. Vogelstein, Y. Qin, B. Cocanougher, K. Eichler, M. Zlatic, et al. Semiparametric spectral modeling of the drosophila connectome. arXiv preprint 1705.03297, 2017.

I. M. Ribeiro, M. Drews, A. Bahl, C. Machacek, A. Borst, and B. J. Dickson. Visual projection neurons mediating directed courtship in drosophila. Cell, 174(3):607–621, 2018.

L. K. Scheffer, C. S. Xu, M. Januszewski, Z. Lu, S.-y. Takemura, K. J. Hayworth, G. B. Huang, K. Shinomiya, J. Maitlin-Shepard, S. Berg, et al. A connectome and analysis of the adult drosophila central brain. elife, 9:e57443, 2020.

P. Schlegel, Y. Yin, A. S. Bates, S. Dorkenwald, K. Eichler, P. Brooks, D. S. Han, M. Gkantia, M. Dos Santos, E. J. Munnelly, et al. Whole-brain annotation and multi-connectome cell typing of drosophila. Nature, 634(8032):139–152, 2024.

J. D. Seelig and V. Jayaraman. Neural dynamics for landmark orientation and angular path integration. Nature, 521(7551):186–191, 2015.

R. Sen, M. Wu, K. Branson, A. Robie, G. M. Rubin, and B. J. Dickson. Moonwalker descending neurons mediate visually evoked retreat in drosophila. Current Biology, 27(5):766–771, 2017.

H. S. Seung. Reading the book of memory: sparse sampling versus dense mapping of connectomes. Neuron, 62(1):17–29, 2009.

R. Tanaka and D. A. Clark. Neural mechanisms to exploit positional geometry for collision avoidance. Current Biology, 32(11):2357–2374, 2022.

D. Turner-Evans, S. Wegener, H. Rouault, R. Franconville, T. Wolff, J. D. Seelig, S. Druckmann, and V. Jayaraman. Angular velocity integration in a fly heading circuit. Elife, 6:e23496, 2017.

A. H. Williams, E. Kunz, S. Kornblith, and S. Linderman. Generalized shape metrics on neural representations. Advances in Neural Information Processing Systems, 34:4738–4750, 2021.

M. Wu, A. Nern, W. R. Williamson, M. M. Morimoto, M. B. Reiser, G. M. Card, and G. M. Rubin. Visual projection neurons in the drosophila lobula link feature detection to distinct behavioral programs. Elife, 5:e21022, 2016.

M. Xu. Understanding graph embedding methods and their applications. SIAM Review, 63(4):825–853, 2021.

D. L. Yamins and J. J. DiCarlo. Using goal-driven deep learning models to understand sensory cortex. Nature neuroscience, 19(3):356–365, 2016.

